# Super food or super toxic? Turmeric and spirulina as culprits for the toxic effects of food dyes in *Drosophila*

**DOI:** 10.1101/2023.10.26.564230

**Authors:** Rebecca von Hellfeld, Craig Christie, Davina Derous, Juliano Morimoto

**Author notes:** Equal contributions (Rebecca von Hellfeld, Juliano Morimoto). **Authors’ contributions** RvH and JM analysed the data and wrote the manuscript. CC, DD, and JM collected the data. All authors revised the manuscript and approved its submission to the journal. **Data availability statement** Data will be available in Dryad upon acceptance of the manuscript.

## Abstract

Prolonged exposure to food dyes, even for those considered safe for consumption, are known to have toxic effects. However, we lack a proper understanding of the underlying compounds that are responsible for the observed toxicity. In this study, we tested the toxic physiological effects of three common commercially available natural food dyes (red, green, blue), and their main ingredients (turmeric and spirulina), on *Drosophila melanogaster* oviposition, larval development, and larval foraging behaviour. Larval development and egg-to-adult survival was significantly impacted by blue and green dyes. These effects were recapitulated when flies were fed with increasing concentrations of turmeric and spirulina, suggesting that turmeric is a toxic component of the food dye. Red dye, which contains neither turmeric or spirulina, had little impact on fly health and behaviour. Green and blue food dyes decreased egg laying, an effect similar to that observed in increasing concentrations of turmeric and, to a lesser extent, spirulina. When given a choice, larvae preferred to feed as follows: control > red > blue > green diet patches, a pattern inversely correlating with the previously observed toxicity. Our results show that, despite turmeric being often considered a super food, it can have toxic effects that the impact organismal physiology and health.

## Introduction

The psychology of colour influences consumer behaviour and has been the target of marketing strategies in various industries, including food and beverages (J. S. Kumar, 2017I). Although food safety standards ensure that such additives are safe for consumption, recent studies have determined that synthetic food dyes are a major source of intoxication, leading to severe health implications (Dey & Nagababu, 2022). In mice, tartrazine (also known as Yellow 5, E102) decreased learning and memory, and in rats, it significantly increased activity in open field tests, likely due to brain damage induced by reactive oxygen species (Gao et al., 2011). Furthermore, a three-generational study determined the adverse effects of tartrazine on various neuro-behavioural parameters in mice (Tanaka et al., 2008). For instance, dye exposure led to accelerated swimming direction development in F1 and F2 males, surface rearing and exploratory behaviour in F1 males were impaired, and olfactory orientation was improved in F2 males. Other food dyes, e.g., Red 2 (E123), have also been implicated in toxicity effects (Mpountoukas et al., 2010). Thus, more studies are urgently needed to understand better the health implications of food dyes and the underlying components used in their production.

Studies in invertebrate models, primarily *Drosophila* flies, have corroborated the results from vertebrate model organisms (Bilder & Irvine, 2017; Link & Bellen, 2020; Reiter et al., 2001). Turmeric, often classified as a super food, increased larval mortality and decreased adult fly survival (Uysal et al., 2015). Moreover, studies have also assessed the toxic impact of synthetic food dyes on *Drosophila* development, linking, e.g., ‘brilliant blue’ and ‘sunset yellow’ exposure to developmental delay, reduced locomotion, morphological changes, and paralysis, with indications of potential interference with neurotransmitters (P. B. Kumar et al., 2020). Likewise, ‘brilliant blue’ significantly decreased *Drosophila* longevity, with other dye colours having similar but less pronounced impacts (Türkoğlu et al., 2015). Population decline was also determined by various yellow and red synthetic dyes in this species (Uysal et al., 2017). Despite growing evidence of toxic effects of food dyes, decades of studies in *Drosophila* have used food dyes to quantify food intake without fully acknowledging the potential toxic effects of these compounds. For instance, Tanimura et al., (1982) utilised red and blue synthetic dyes to determine the preference of *Drosophila* strains for different types of sugar, but no preliminary assessment of dye toxicity appears to have been conducted. Similarly, Shell et al., (2018) and Shell & Grotewiel, (2022) highlighted the benefits of using synthetic food dye for assessing fly consumption-excretion, providing the means to detect metabolic activity by measuring the absorbance of excreted dye in a spectrophotometer. This technique formed the basis for further studies investigating food preferences using different synthetic dyes as labels (Mack & Zhang, 2021; Shell et al., 2021). Dyes have also been used to determine how intestinal barrier dysfunction may link to age markers like inflammation and metabolic changes (Rera et al., 2012, 2013), but with no preliminary assessment of the negative impacts of the dyes on development or survival. Therefore, toxicity studies of food dye in *Drosophila* can shed light on the potential effects for human consumption and help improve current protocols in the broader field of nutritional sciences using invertebrates as models.

In this study, we investigated the toxic effects of three commercial food dyes (red, blue, and green) and their main colour-giving compounds (turmeric and spirulina) on female oviposition, larval development, larval foraging behaviour and adult sex ratio in *D. melanogaster*. We conducted three experiments to reveal the toxicity effects of these compounds. In experiment one (‘no choice – dye diet’), adult females were placed in diets made with one of the food dyes and allowed to oviposit freely. We then measured total number of eggs laid as well as egg-to-adult development. In experiment two (‘choice experiment – dye diet’), eggs were placed in foraging arenas (see Morimoto, 2019; Morimoto et al., 2018, 2019) and allowed to forage freely among patches with control diet and dyes, upon which patch utilisation and pupation site were recorded. These two experiments allowed us to determine the toxicity effects of each food dye and the perceived preference for each food dye by individuals. Then, in experiment three (‘no choice – turmeric or spirulina diet’), we used the same approach as experiment one but placed females in diets with increasing concentrations of either turmeric or spirulina to determine potential toxic effects. Based on the manufacturer’s ingredient list, we expected the red dye to have no effect on any of the traits measured, while we expected both blue and green dyes to negatively affect the traits due to the addition of turmeric in their recipe. However, we had no *a priori* information as to the proportions of turmeric and spirulina added to the blue and green dyes, and, therefore, had no *a priori* predictions as to which dye should have the strongest toxic effects. Our findings question turmeric status as a superfood by showing that it can be highly toxic to a developing model organism.

## Material and Methods

### Fly stock and egg collection

We used an outbred population of *Drosophila melanogaster* from Brittany, France. Fly stocks were maintained in large populations (>1,000 individuals) at 20 °C, approximately 40 % humidity, with a 12:12 h light:dark photoperiod, and fed a yeast-sucrose diet (‘control’ diet) as described in Morimoto, (2022). Eggs were collected by placing two bottles with the control diet in the stock population cage for 24 h. Bottles were retrieved and maintained at 20 °C until adult emergence. Within 6 h of adult emergence, five randomly selected virgin males and females were placed in vials with 5 mL of diets containing food dyes (experiment one), turmeric or spirulina (experiment three). Twenty vials with control diets were reserved, and the groups were allowed to oviposit for 15 h, after which eggs were then collected from the vials and immediately placed into the foraging arenas (experiment two). All experiments were conducted at 20 °C with 24:0 h light:dark photoperiod.

### Statistical analysis

All statistical analyses were conducted in RStudio 2023.06.0+421 ‘Mountain Hydrangea’. We used the following packages: ‘dplyr’ (Wickham, François, et al., 2023); ‘tidyr’ (Wickham, Vaughan, et al., 2023); ‘lmerTest’ (Kuznetsova et al., 2020); ‘lmer4’ package (Bates et al., 2015), ‘car’ package (Fox & Weisberg, 2019), ‘nnet’ (Ripley & Venables, 2023); ‘stringr’ (Wickham, 2022); and ‘agricolae’ (de Mendiburu, 2023). Statistical models for each experiment are given in the sections below. All figures were created with ‘ggplot2’ (Wichkam, 2016) and ‘patchwork’ (Pederson, 2023). Complete model outputs are given in Table S1.

### Experimental design

#### Experiment one: No choice – dye diets

Flies were added to one of four diet treatments (*N*_*total*_ *=* 16): either control (no dye) or diets with 20 % of one of three commercial natural food dyes: blue, green, and red (Betty Winters®). The concentration was selected to determine potential impacts of diet without reducing macronutrient intake to an extent that would elicit negative impacts on development or behaviour. The concentration used here exceeds those utilised in previous studies (see e.g., Tanimura et al., 1982), as the solid food used here is not as colourless as liquid fly diet, thus needing more dye to become visible in uptake. As food was not limited in any of the exposure experiments, it was not necessary to replace 20 % of the control diet with water to account for potential differences in macronutrients.

Adults were allowed to interact freely, mate, and females were allowed to lay eggs for 48 h, after which adults were discarded. The eggs were then counted using a Leica M9i stereomicroscope and maintained as described above to complete development until adult emergence. Pupation was scored twice daily (mornings and afternoons) by counting the number of pupae per food patch, as a measure of developmental time (egg-to-pupa). After pupation was first observed, all vials were maintained for 10 days, allowing adults to emerge before being frozen at -20 °C for 48 h. Where larvae did not pupate within the specified 10 days, additional 20 days were given. If pupation did not occur after 30 days, the experiment was terminated (‘censored’).. The number of emerged males and females in each diet treatment were compared to determine the sex ratio of emerging adults across diet treatments. Egg-to-adult viability was calculated as the number of adults divided by the number of eggs in the vial x 100. We fitted a generalised mixed model (GLMER) with egg batch as the random variable and food dye as the independent variable. To analyse the counts of the number of eggs laid in each diet, we fitted a model with Poisson error distribution. For developmental time and sex ratio, we used a Gamma error distribution to best fit the data. For the egg-to-adult viability percentage, we used a Binomial error distribution. P-values were obtained using the ANOVA function of the ‘car’ package and the pairwise comparisons from the ‘summary’ function in base R.

#### Experiment Two: Choice – dye diets

30 or 60 eggs were placed at the centre of a foraging arena (*N*_*total*_ *= 8* per egg density), which was designed as in Morimoto, (2019); Morimoto et al., (2018, 2019). Briefly, a 90mm petri dish was covered with a solution of 2 % agar, and 4 equally spaced discs were removed to make space for discs representing one of the four diets (control, red, blue, and green). Discs were made using a 15 mL Falcon tube. The two egg densities were used because previous studies have shown that social group density influences accuracy and speed of larval foraging decisions (Morimoto et al., 2018, 2019). Larvae had free choice of foraging patch and pupation site. Foraging arenas were frozen at -20 °C after 11 days of incubation, after which pupation site was scored by counting the number of pupae in the proximity of each of the diet patches. To estimate food utilisation, we photographed each diet patch in each replicate, and created a score of food utilisation that ranged from 0 to 3 (0 = not utilised, 1 = poorly utilised, 2 = utilised, 3 = heavily utilised; see Figure S1 for scoring criteria). Three independent scorers (JM, DD, CC) analysed the images and assigned scores. The average of the scores were used as food utilisation index (see below). We fitted a linear mixed model with batch as random factor to investigate pupation number in relation to food dye, egg density (as factor with two levels) and the interaction between dye and egg density. We then fitted a multinomial regression to model the probability of pupation in each of the sites, and 95 % confidence intervals of estimates were obtained using the ‘confint’ function in R.

#### Experiment three: no choice – turmeric or spirulina diets

We adopted the same experimental design as described for experiment one, but here the diets had increasing concentrations (0 %, 1 %, 2.5 %, 5 %, and 10 % w/v) of either pesticide-free turmeric or pesticide-free spirulina (Sevenhills Wholefoods®). These concentrations were selected to determine potential toxic effects, as well as understanding concentration-related impacts. Each compound was analysed separately (*N*_*total*_ = 30 per compound). To establish a baseline for comparison with the food dye experiment, we also had diet treatments containing each of the food dyes tested in experiment 1 (*N* = 3 per food dye) at 20 % v/v concentration. We fitted a general linear model with log-transformed number of eggs laid, developmental time, and untransformed sex ratio, with turmeric or spirulina concentration as the independent variable. For the proportion of adult emergence, we fitted a generalised linear model with Binomial error and quasi extension to account for the overdispersion of the data. P-values were obtained from F-statistics in the ‘anova’ function in R.

## Results

### Experiment one: Food dye significantly affected fly development

Food dye affected female oviposition behaviour (χ^2^ = 47.833; *degrees of freedom (df)* = 3; *p* <0.001). Compared to the control diet, females laid a similar number of eggs in the green dye diet (*estimate* ± *standard error:* -0.005 ± 0.050; *p* = 0.919). Significantly fewer eggs were laid in diets with the blue dye (0.144 ± 0.052; *p* = 0.006) and significantly more eggs in diets with red dye (0.194 ± 0.048; *p* <0.001; Fig 1a). Developmental time (egg-to-adult) was also significantly impacted by the food dye (χ^2^ = 345.67; *df* = 3; *p* <0.001). Compared to the control diet, larval developmental time was significantly longer in both the blue (-0.009 ± 0.003; *p* <0.001) and green dye diets (-0.034 ± 0.002; *p* <0.001). In red dye diets, larval developmental time was weakly but statistically significantly faster than controls (0.005 ± 0.003, *p* = 0.049; Fig 1b). Egg-to-adult viability was also significantly affected by food dye (χ^2^ = 179.37; *df* = 3; *p* <0.001) largely driven by green dye almost completely abolished fly viability (-1.295 ± 0.104; *p* <0.001; Fig 1c). There was no statistically significant effect of food dye on adult sex ratio (χ^2^ = 4.398; *df* = 3; *p* = 0.221; Fig 1d).

**Figure 1.**
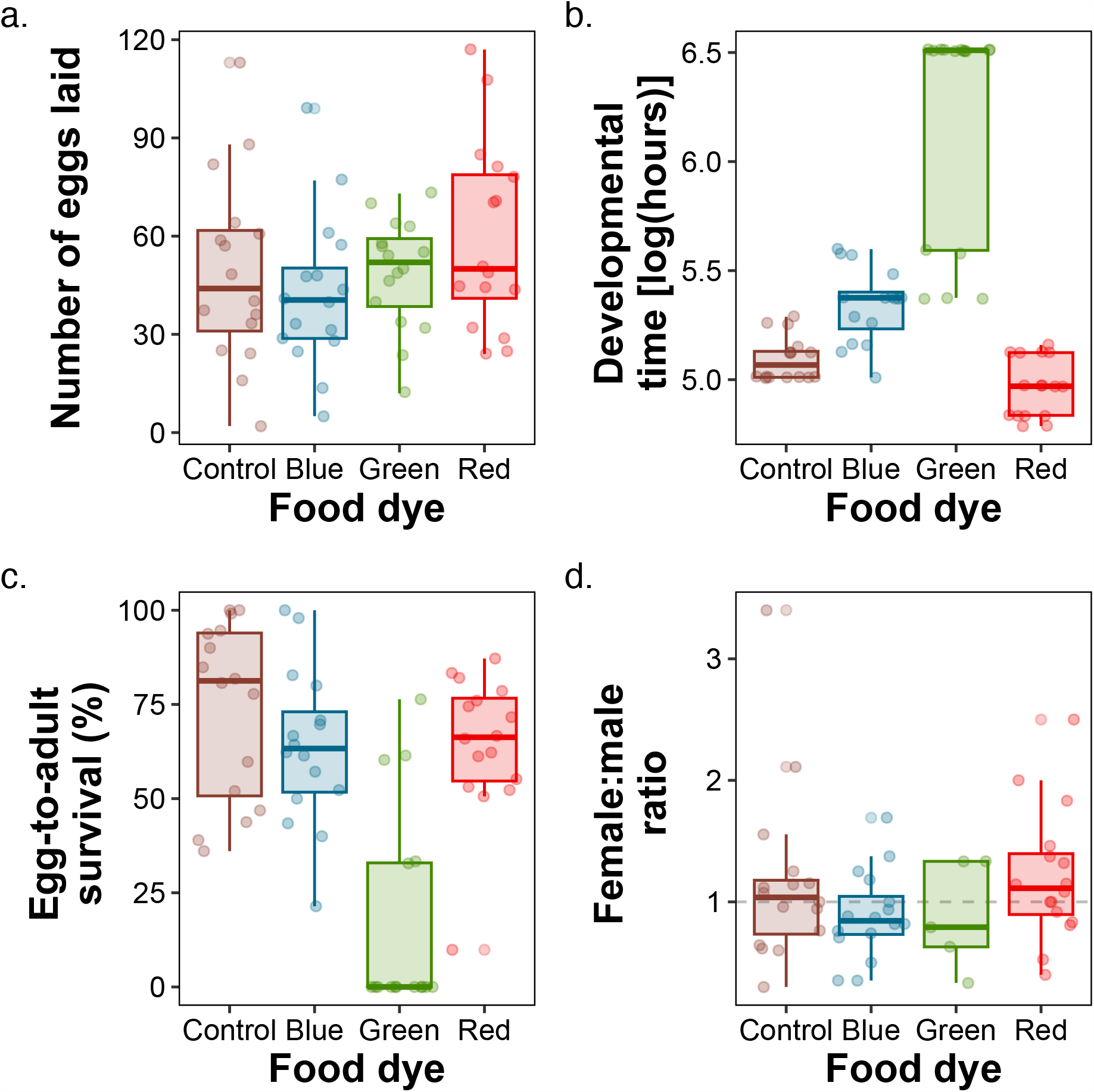
Food dye effects on female oviposition and larval development. (a) the number of eggs laid, (b) the developmental time log(hours) from egg to adult, (c) egg-to-adult survival ratio, and (d) adult sex ratio, where dashed line indicates a 1:1 male:female ratio.

### Experiment 2: Larvae avoid utilising dye-containing diets

Next, we analysed how flies performed when given a choice among food patches containing different food dyes. There was a statistically significant effect of food dye on food use index (F_3,254_ = 10.337; *p* <0.001), but no effect of egg density (F_1,254.01_ = 0.647; *p* = 0.421) or the interaction between egg density and food dye (F_3,254_ = 0.653; *p* = 0.581; Fig 2a). Flies tended to prefer control diets, followed by red, blue, and green, in reverse order of the toxic effects observed for development (Fig 2a). However, there was a higher probability of pupation in green, blue, red and control in smaller groups (egg density = 30) as opposed to larger groups (egg density = 60; χ^2^ = 88.872; *df* = 3, *p* <0.001; Fig 2b). There was no effect of day (χ^2^ = 0.484; *df* = 3; *p* = 0.922) or the interaction of day and egg density (χ^2^ = 0.479; *<u>df</u>* = 3; *p* = 0.923) on the probability of pupation. There was a statistically significant interaction between egg density and pupation number (F_3,183_ = 5.162; *p* = 0.001), driven by the increase in pupation number in all food dye patches from days 9 to 11 in larger groups (egg density = 60) but a stable (red and control), or only marginal increase (blue and green) in pupation number in the food patch in smaller groups (egg density = 30; Fig 2c).

**Figure 2.**
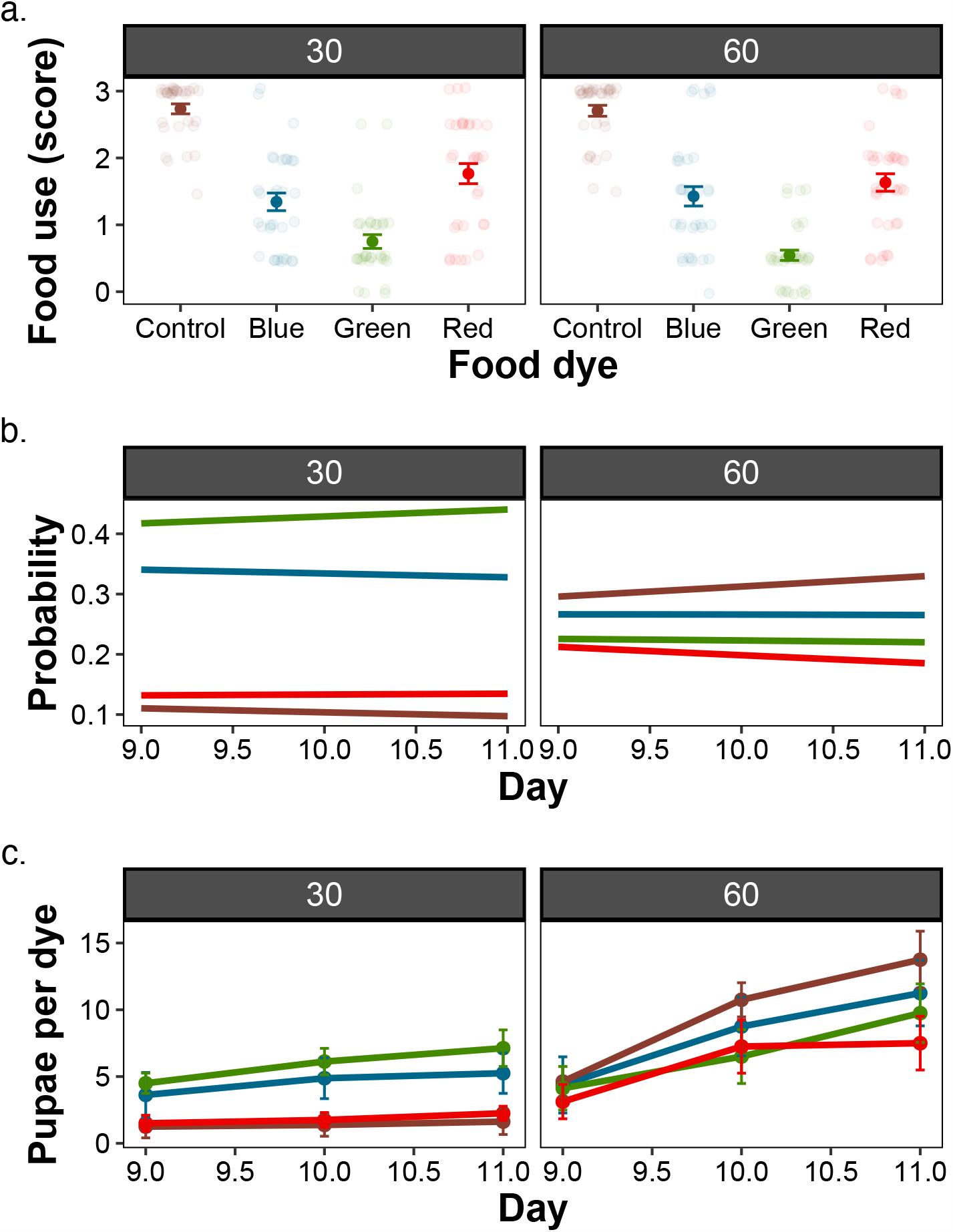
Food dye effects on food utilisation and pupation site. (a) the use of substrate, (b) the probability of pupation per substrate, and (c) the number of pupae per dye for egg-cohort sizes of 30 and 60 eggs.

### Experiment 3: Turmeric is the primary developmental toxicant in food dye

Lastly, we investigated which compound was the toxicant in green and blue food dyes. According to the manufacturer, these dyes were a mixture of turmeric and spirulina. We, therefore, tested the effects of increasing concentration of these compounds on fly development. Increasing turmeric concentration decreased egg laying (F_4,25_ = 24.78; *p* <0.001), extended developmental time (F_4,25_ = 1251.4; *p* <0.001), and decreased egg to adult viability (F_4,25_ = 76.32; *p* <0.001), effectively recapitulating the toxic effects observed in the green food dye for a concentration of 10 % turmeric, and the blue dye for a concentration of 2.5 % turmeric (Fig 3a-c). Increasing spirulina concentration extended developmental time (F_4,25_ = 4.442; *p* = 0.008) and decreased egg to adult viability (F_4,25_ = 2.994; *p* = 0.037). There was no effect of spirulina concentration on egg laying (F_4,25_ = 2.247; *p* = 0.092). Neither was the adult sex ratio affected by turmeric (F_3,21_ = 0.5677; *p* = 0.642) or spirulina (F_4,25_ = 0.340, *p* = 0.847), although the highest concentration of turmeric had no adult emergence (Fig 3a-d).

**Figure 3.**
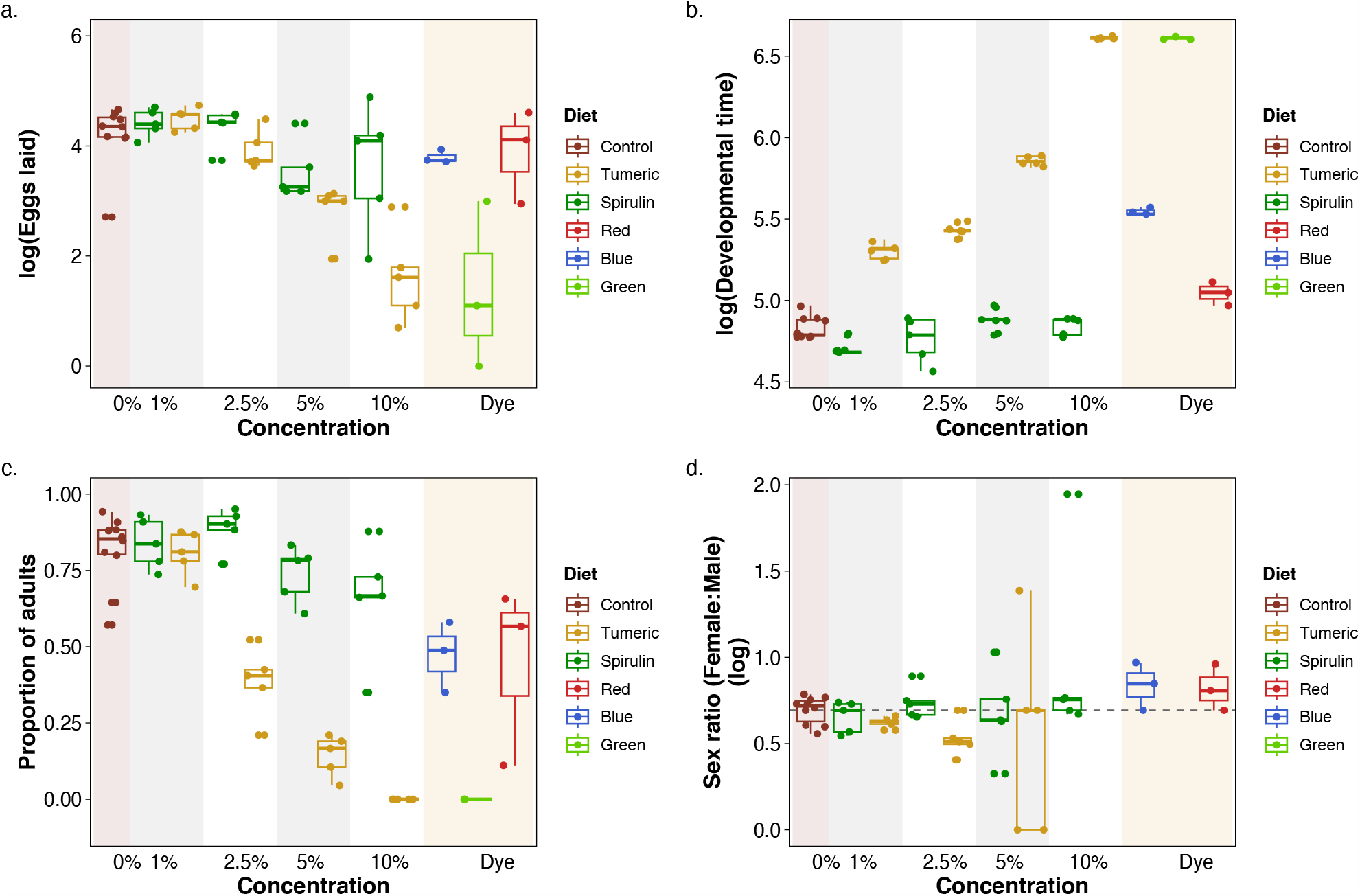
Tumeric and spirulina affect female oviposition and larval development. (a) the number of eggs laid, (b) the developmental time log(hours) from egg to adult, (c) egg-to-adult survival ratio, and (d) adult sex ratio, with dashed line indicating a 1:1 male:female ratio .

## Discussion

Food dyes are commonly used for human consumption or as visual markers for experiments in the laboratory involving food intake (see e.g., Shell et al., 2018, 2021; Shell & Grotewiel, 2022). Here, we showed that food dyes can display toxic effects, using *Drosophila* as a model organism. We found that green and blue dyes were toxic to the developing fly. Through follow-on studies, we found that turmeric and, to a lesser extent, spirulina were likely the toxic agents responsible for the observed differences. Increasing the concentration of turmeric almost perfectly recapitulated the adverse effects observed for oviposition, developmental time, and egg-to-adult viability observed in green and blue dyes (Fig 3). These results confirm previous findings in flies that turmeric can be highly toxic and decrease survival in both larvae and adults (Uysal et al., 2015). Importantly, our findings show that experimental protocols that use food dyes to measure food intake must be mindful of the dyes’ potential side effects on larval food intake and life-histories.

A previous experiment by Kumar et al., (2020) showed that exposure to 20-30 % of ‘Brilliant Blue FCF’ dye decreased hatching success, pupation, and adult eclosion in *Drosophila*. This dye was chemically manufactured, and thus, the toxicity effects are likely due to toxic chemical elements as opposed to natural compounds tested here. Marcus et al., (2018) showed that female *Drosophila* preferred to lay eggs on green over blue and aqua-coloured sheets placed under clear agar, highlighting the importance of colour itself for egg allocation and egg-laying for the species. We did not observe this effect with a higher incidence of laying in the control and red dye diets when compared with flies housed with green and blue diets. This effect is probably because of some gustatory and/or olfactory cues from the turmeric and spirulina present into those dyes, which females used for oviposition selection.

A study in *Drosophila* found that increasing turmeric concentration reduced adult survival (Uysal et al., 2015), while another found that turmeric had no impact on oviposition of this species (Rawal et al., 2014). Only one study examined the impact of curcumin, the main active compound in turmeric, on *Drosophila* larval development, reporting increased development times but without further investigations into the underlying molecular mechanisms (Soh et al., 2013). Other studies suggested that curcumin intake induces oxidative stress response, causing developmental impairments (Esquivel et al., 2020; Puri et al., 2021; Shen et al., 2013). Moreover, curcuminoids are known to act as photoactivatable insecticides, wherein the combination of oxygen and light with their photosensitiser property trigger photochemical reactions culminating in the destruction of e.g., membranes, mitochondria, and proteins (Ben Amor & Jori, 2000) – which is known as photodynamic inactivation (PDI). This phenomenon has been linked to the formation of reactive oxygen species (ROS) (Allison et al., 2008; Huang et al., 2009; Smijs et al., 2007). We hypothesise that curcumin is the main driver of the effect of turmeric observed in our data, and further studies focusing on this compound are being conducted. Curcumin has also been found to reduce neurotoxicity in transgenic flies that are models for Alzheimer’s disease, suggesting that curcumin and, perhaps, turmeric may have tissuespecific effects (Caesar et al., 2012). Studies in another fly species (*Bactrocera cucurbitae*) found that developmental time was significantly increased with curcumin exposure, which they attributed to an aversion to the diet due to post-ingestion toxic effects (Puri et al., 2021). This aversion directly affected the weight needed for pupation (Davidowitz et al., 2003), delaying overall growth. Puri et al., (2021) further determined that *B cucurbitae* egg-to-adult survival decreased in a concentration-dependent manner by curcumin.

A recent study in western honeybees *Apis mellifera* showed that the synthetic blue dye ‘Brilliant Blue FCF’ significantly reduced larval survival compared to the control diet (Ehrenberg et al., 2019). They hypothesised that the blue dye could either elicit a change in nutrient uptake, be metabolised differently to other dyes, leading to complex intermediates, or affect metamorphosis and pupal development through impacts on resource allocation. Our study did not allow us to investigate the underlying mechanisms, but it is likely that both turmeric and, to a smaller extent spirulina, modulate excretion pathways that are associated with toxicity, such as the uric acid metabolism pathway. Recently, we have shown that the uric acid metabolism is a key modulator of diet- and toxic-dependent responses in *Drosophila* (Morimoto, 2022). Thus, it is likely that the toxic effects observed here are, at least partly, related to this pathway. Future studies using molecular tools will allow us to characterise this and help gain insight into the molecular routes through which turmeric and spirulina realise their toxic effects.

Previous studies investigated the benefits of rearing animals intended as fish feed on spirulina (Ratti et al., 2023; Tu et al., 2022). Qiu et al., (2019) determined that higher concentrations of microalgae in diets for *Drosophila* significantly decreased egg-to-adult viability, whilst also leading to lower adult body weight. Qiu et al., (2019) also assessed the macronutrients of the algal mixture, which had a protein:carbohydrate ratio of 44:26 (1.69:1), as well as containing more than 25 % lipids, whilst standard fly food is more than 50 % carbohydrates and contains no lipids. It is known that the optimum diet for *Drosophila* body size is a protein-to-carbohydrate ratio of 1.5:1, which is comparable to that of spirulina(Rodrigues et al., 2015). Moreover, increasing lipid concentration in the diet appears to have no effect on fly development and body weight (Reis, 2016). Therefore, diet macronutrient composition alone cannot fully explain the results reported by Qiu et al., (2019). Recently, we showed that diet affects toxicity, and thus, there may be combinations of nutrients and toxic compounds that potentialise the toxicity of diet compounds (e.g., see Morimoto, 2022). Nevertheless, our results show that spirulina, although relatively less toxic than turmeric, can still induce developmental delays and decrease egg to adult viability when in high concentrations.

## Conclusions

Turmeric and spirulina are often considered to be superfoods. Here, we show that they can also be super toxic in the *Drosophila* model. As a result, products reliant on these compounds, such as food dyes, may prove toxic to flies and potentially, higher animals. Future studies should focus on the molecular pathways through which turmeric and spirulina induce toxicity in flies, which can reveal candidate genes for toxicity effects in vertebrates and establish the fly as a model for identification and isolation of toxic compounds. Our findings in the invertebrate model serve as a cautionary tale for the indiscriminate labelling of foods as causing no harm when consumed.

## Acknowledgements

JM is funded by BBSRC grant (BB/V015249/1). RvH is funded through DEFRA project (ETPP-33/C10).

## Authors’ contributions

CC, DD, and JM collected the data. RvH and JM analysed the data and wrote the manuscript. All authors revised the manuscript and approved its submission to the journal.

## Data availability statement

Data will be available in Dryad upon acceptance of the manuscript.

## Supplementary Material

**Figure S1.** Example of scoring criteria

## References

Allison, R. R., Mota, H. C., Bagnato, V. S., & Sibata, C. H. (2008). Bio-nanotechnology and photodynamic therapy—State of the art review. Photodiagnosis and Photodynamic Therapy, 5(1), 19–28. 10.1016/j.pdpdt.2008.02.001

Bates, D., Mächler, M., Bolker, B., & Walker, S. (2015). Fitting Linear Mixed-Effects Models Using lme4. Journal of Statistical Software, 67(1). 10.18637/jss.v067.i01

Ben Amor, T., & Jori, G. (2000). Sunlight-activated insecticides: historical background and mechanisms of phototoxic activity. Insect Biochemistry and Molecular Biology, 30(10), 915–925. 10.1016/S0965-1748(00)00072-2

Bilder, D., & Irvine, K. D. (2017). Taking stock of the Drosophila research ecosystem. Genetics, 206(3), 1227–1236. 10.1534/genetics.117.202390

Caesar, I., Jonson, M., Nilsson, K. P. R., Thor, S., & Hammarström, P. (2012). Curcumin promotes A-beta fibrillation and reduces neurotoxicity in transgenic Drosophila. PLoS ONE, 7(2), e31424. 10.1371/journal.pone.0031424

Davidowitz, G., D’Amico, L. J., & Nijhout, H. F. (2003). Critical weight in the development of insect body size. Evolution and Development, 5(2), 188–197. 10.1046/j.1525-142X.2003.03026.x

de Mendiburu, F. (2023). agricolae: Statistical Procedures for Agricultural Research. https://cran.r-project.org/package=agricolae

Dey, S., & Nagababu, B. H. (2022). Applications of food color and bio-preservatives in the food and its effect on the human health. Food Chemistry Advances, 1, 100019. 10.1016/j.focha.2022.100019

Ehrenberg, S., Lewkowski, O., & Erler, S. (2019). Dyeing but not dying: Colourful dyes as a non-lethal method of food labelling for in vitro-reared honey bee (Apis mellifera) larvae. Journal of Insect Physiology, 113, 1–8. 10.1016/j.jinsphys.2018.12.008

Esquivel, A. R., Douglas, J. C., Loughran, R. M., Rezendes, T. E., Reed, K. R., Cains, T. H. L., Emsley, S. A., Paddock, W. A., Videau, P., Koyack, M. J., & Paddock, B. E. (2020). Assessing the influence of curcumin in sex specific oxidative stress, survival, and behavior in Drosophila melanogaster. Journal of Experimental Biology. 10.1242/jeb.223867

Fox, J., & Weisberg, S. (2019). An R Companion to Applied Regression (3rd ed.). https://socialsciences.mcmaster.ca/jfox/Books/Companion/

Gao, Y., Li, C., shen, J., Yin, H., An, X., & Jin, H. (2011). Effect of food azo dye tartrazine on learning and memory functions in mice and rats, and the possible mechanisms involved. Journal of Food Science, 76(6), T125–T129. 10.1111/j.1750-3841.2011.02267.x

Huang, Y.-Y., Chen, A. C.-H., Carroll, J. D., & Hamblin, M. R. (2009). Biphasic dose response in low level light therapy. Dose-Response, 7(4), dose-response.0. 10.2203/dose-response.09-027.Hamblin

Kumar, J. S. (2017). The psychology of colour influences consumers’ buying behaviour – A diagnostic study. Ushus - Journal of Business Management, 16(4), 1–13. 10.12725/ujbm.41.1

Kumar, P. B., Moses, S., Arunkumar, B., & Ramamurthy, D. S. (2020). Effect of developmental toxicity in Drosophila melanogsater on exposure to different food dyes (Brilliant Blue and Sunset Yellow). International Journal of Science and Research, 9(12). https://www.ijsr.net/archive/v9i12/SR201212195534.pdf

Kuznetsova, A., Bruun Brockhoff, P., Bojesen Christensen, R. H., & Podenphant Jensen, S. (2020). lmerTest: Tests in Linear Mixed Effect Models. https://CRAN.R-pro-ject.org/package=lmerTest

Link, N., & Bellen, H. J. (2020). Using Drosophila to drive the diagnosis and understand the mechanisms of rare human diseases. Development, 147(21). 10.1242/dev.191411

Mack, J. O., & Zhang, Y. V. (2021). A rapid food-preference assay in Drosophila. Journal of Visualized Experiments, 168. 10.3791/62051

Marcus, M., Burnham, T. C., Stephens, D. W., & Dunlap, A. S. (2018). Experimental evolution of color preference for oviposition in Drosophila melanogaster. Journal of Bioeconomics, 20(1), 125–140. 10.1007/s10818-017-9261-z

Morimoto, J. (2019). Foraging decisions as multi-armed bandit problems: Applying reinforcement learning algorithms to foraging data. Journal of Theoretical Biology, 467, 48–56. 10.1016/j.jtbi.2019.02.002

Morimoto, J. (2022). Uric acid metabolism modulates diet-dependent responses to intraspecific competition in Drosophila larvae. IScience, 25(12), 105598. 10.1016/j.isci.2022.105598

Morimoto, J., Nguyen, B., Tabrizi, S. T., Ponton, F., & Taylor, P. (2018). Social and nutritional factors shape larval aggregation, foraging, and body mass in a polyphagous fly. Scientific Reports, 8(1), 14750. 10.1038/s41598-018-32930-0

Morimoto, J., Tabrizi, S. T., Lundback, I., Mainali, B., Taylor, P. W., & Ponton, F. (2019). Larval foraging decisions in competitive heterogeneous environments accommodate di-ets that support egg-to-adult development in a polyphagous fly. Royal Society Open Science, 6(4). 10.1098/rsos.190090

Mpountoukas, P., Pantazaki, A., Kostareli, E., Christodoulou, P., Kareli, D., Poliliou, S., Mourelatos, C., Lambropoulou, V., & Lialiaris, T. (2010). Cytogenetic evaluation and DNA interaction studies of the food colorants amaranth, erythrosine and tartrazine. Food and Chemical Toxicology, 48(10), 2934–2944. 10.1016/j.fct.2010.07.030

Pederson, T. L. (2023). patchwork: The Composer of Plots. https://patchwork.data-imaginist.com, https://github.com/thomasp85/patchwork

Puri, S., Singh, S., & Sohal, S. K. (2021). Growth retarding effect of curcumin on Bactrocera cucurbitae (Coquillett) larvae. Archives of Phytopathology and Plant Protection, 54(13– 14), 722–735. 10.1080/03235408.2020.1857134

Qiu, S., Wang, S., Xiao, C., & Ge, S. (2019). Assessment of microalgae as a new feeding additive for fruit fly Drosophila melanogaster. Science of The Total Environment, 667, 455–463. 10.1016/j.scitotenv.2019.02.414

Ratti, S., Zarantoniello, M., Chemello, G., Giammarino, M., Palermo, F. A., Cocci, P., Mosconi, G., Tignani, M. V., Pascon, G., Cardinaletti, G., Pacetti, D., Nartea, A., Parisi, G., Riolo, P., Belloni, A., & Olivotto, I. (2023). Spirulina-enriched substrate to rear black soldier fly (Hermetia illucens) prepupae as alternative aquafeed ingredient for rainbow trout (Oncorhynchus mykiss) diets: Possible effects on zootechnical performances, gut and liver health status, and fillet quality. Animals, 13(1), 173. 10.3390/ani13010173

Rawal, S., Singh, P., Gupta, A., & Mohanty, S. (2014). Dietary Intake of Curcuma longa and Emblica officinalis Increases Life Span in Drosophila melanogaster. BioMed Research International, 2014, 1–7. 10.1155/2014/910290

Reis, T. (2016). Effects of synthetic diets enriched in specific nutrients on Drosophila development, body fat, and lifespan. PLOS ONE, 11(1), e0146758. 10.1371/journal.pone.0146758

Reiter, L. T., Potocki, L., Chien, S., Gribskov, M., & Bier, E. (2001). A systematic analysis of human disease-associated gene sequences In Drosophila melanogaster. Genome Research, 11(6), 1114–1125. 10.1101/gr.169101

Rera, M., Azizi, M. J., & Walker, D. W. (2013). Organ-specific mediation of lifespan extension: More than a gut feeling? Ageing Research Reviews, 12(1), 436–444. 10.1016/j.arr.2012.05.003

Rera, M., Clark, R. I., & Walker, D. W. (2012). Intestinal barrier dysfunction links metabolic and inflammatory markers of aging to death in Drosophila. Proceedings of the National Academy of Sciences, 109(52), 21528–21533. 10.1073/pnas.1215849110

Ripley, B., & Venables, W. (2023). nnet: Feed-Forward Neural Networks and Multinomial Log-Linear Models. https://cran.r-project.org/package=nnet

Rodrigues, M. A., Martins, N. E., Balancé, L. F., Broom, L. N., Dias, A. J. S., Fernandes, A. S. D., Rodrigues, F., Sucena, É., & Mirth, C. K. (2015). Drosophila melanogaster larvae make nutritional choices that minimize developmental time. Journal of Insect Physiology, 81, 69–80. 10.1016/j.jinsphys.2015.07.002

Shell, B. C., & Grotewiel, M. (2022). Identification of additional dye tracers for measuring solid food intake and food preference via consumption-excretion in Drosophila. Scientific Reports, 12(1). 10.1038/s41598-022-10252-6

Shell, B. C., Luo, Y., Pletcher, S., & Grotewiel, M. (2021). Expansion and application of dye tracers for measuring solid food intake and food preference in Drosophila. Scientific Reports, 11(1). 10.1038/s41598-021-99483-7

Shell, B. C., Schmitt, R. E., Lee, K. M., Johnson, J. C., Chung, B. Y., Pletcher, S. D., & Grotewiel, M. (2018). Measurement of solid food intake in Drosophila via consumptionexcretion of a dye tracer. Scientific Reports, 8(1). 10.1038/s41598-018-29813-9

Shen, L.-R., Parnell, L. D., Ordovas, J. M., & Lai, C.-Q. (2013). Curcumin and aging. Bio-Factors, 39(1), 133–140. 10.1002/biof.1086

Smijs, T. G. M., Bouwstra, J. A., Schuitmaker, H. J., Talebi, M., & Pavel, S. (2007). A novel ex vivo skin model to study the susceptibility of the dermatophyte Trichophyton rubrum to photodynamic treatment in different growth phases. Journal of Antimicrobial Chemotherapy, 59(3), 433–440. 10.1093/jac/dkl490

Soh, J.-W., Marowsky, N., Nichols, T. J., Rahman, A. M., Miah, T., Sarao, P., Khasawneh, R., Unnikrishnan, A., Heydari, A. R., Silver, R. B., & Arking, R. (2013). Curcumin is an early-acting stage-specific inducer of extended functional longevity in Drosophila. Experimental Gerontology, 48(2), 229–239. 10.1016/j.exger.2012.09.007

Tanaka, T., Takahashi, O., Oishi, S., & Ogata, A. (2008). Effects of tartrazine on exploratory behavior in a three-generation toxicity study in mice. Reproductive Toxicology, 26(2), 156–163. 10.1016/j.reprotox.2008.07.001

Tanimura, T., Isono, K., Takamura, T., & Shimada, I. (1982). Genetic dimorphism in the taste sensitivity to trehalose in Drosophila melanogaster. Journal of Comparative Physiology ? A, 147(4), 433–437. 10.1007/BF00612007

Tu, N. P. C., Ha, N. N., Linh, N. T. T., & Tri, N. N. (2022). Effect of astaxanthin and spirulina levels in black soldier fly larvae meal-based diets on growth performance and skin pigmentation in discus fish, Symphysodon sp. Aquaculture, 553, 738048. 10.1016/j.aquaculture.2022.738048

Türkoğlu, Ş., Benli, D., & Şahin, N. (2015). The effects of five food dyes on the longevity of Drosophila melanogaster. Fresenius Environmetal Bulletin, 24(9). https://www.researchgate.net/publication/286872980

Uysal, H., Genic, S., & Ayar, A. (2017). Toxic effects of chronic feeding with food flazo dyes on Drosophila melanogaster Oregon R. Scientia Iranica, 24(6), 3081–3086. 10.24200/sci.2017.4523

Uysal, H., Semerdöken, S., Çolak, D. A., & Ayar, A. (2015). The hazardous effects of three natural food dyes on developmental stages and longevity of Drosophila melanogaster. Toxicology and Industrial Health, 31(7), 624–629. 10.1177/0748233713480206

Wichkam, H. (2016). ggplot2: Elegant Graphics for Data Analysis. Springer-Verlag New York. https://ggplot2.tidyverse.org.

Wickham, H. (2022). stringr: Simple, Consistent Wrappers for Common String Operations. https://stringr.tidyverse.org, https://github.com/tidyverse/stringr

Wickham, H., François, R., Henry, L., Müller, K., & Vaughan, D. (2023). dplyr: A grammar of data manipulation. https://CRAN.R-project.org/package=dplyr

Wickham, H., Vaughan, D., & Girlich, M. (2023). tidyr: Tidy Messy Data. https://github.com/tidyverse/tidyr

